# Determination of mitochondrial matrix pH during ischemia-reperfusion injury in the isolated mouse heart using a genetically encoded fluorescent sensor

**DOI:** 10.64898/2026.11.14.766482

**Authors:** Abigail V. Giles, Junhui Sun, Barbara Roman, Michael P. Murphy, Robert S. Balaban, Elizabeth Murphy

## Abstract

Mitochondrial matrix pH (pH_mtx)_ regulates cellular energetics, metabolite flux, and ion transport. Despite its importance in cellular energy metabolism and disease, pH_mtx r_emains difficult to measure in the intact heart because pH-sensitive dyes are challenging to selectively target to the mitochondrial matrix. Changes in intracellular pH during cardiac ischemia-reperfusion (I/R) are significant and understanding pH_mtx d_ynamics I/R would provide insight into mechanisms of injury. To overcome uncertainties associated with intracellular dye distribution, the pH-sensitive green fluorescent protein SypHer3 was stably expressed in the mitochondrial matrix of the adult mouse heart using adeno-associated virus (AAV). The isolated perfused heart was subjected to global no-flow ischemia, followed by reperfusion and SypHer3 fluorescence measured using an optical integrating sphere. Calibration of SypHer3 fluorescence fold-change against pH in isolated mitochondria enabled pH_mtx q_uantification from SypHer3 fluorescence in the intact heart, based on an assigned starting pH of 7.47. We found that pH_mtx d_ecreased from 7.45 ± 0.01 (n=5) during normoxia to 6.61 ± 0.20 (n=5) after 20 minutes of ischemia, and reached 7.16 ± 0.01 (n=5) within 1 minute of reperfusion. In paired experiments, we measured relative changes in mitochondrial Ca^2+^ (Ca^2+^_m)_ in the isolated heart during I/R using the genetically encoded, mitochondrial-targeted Ca^2+^ sensor R-GECO1. Because R-GECO1 fluorescence is pH-sensitive, quantification of pH_mtx i_s required to correct R-GECO1 fluorescence for pH-dependent effects. With correction for pH, R-GECO1 fluorescence increased 1.77 ± 0.40-fold (n=7) within 20 minutes of ischemia and returned to baseline within the first minute of reperfusion, consistent with an increase in Ca^2+^_m d_uring ischemia that reversed upon reperfusion. In summary, we developed a fluorescence approach for measuring pH_mtx,_ a key determinant of mitochondrial energetics and ion transport, in the isolated perfused mouse heart, providing time resolved quantitative estimates of pH_mtx d_uring I/R. Furthermore, this work expands the application of fluorescent sensors for interrogating mitochondrial physiology in the intact heart by providing a framework for accounting for pH-dependent effects on probe fluorescence.

## Introduction

Mitochondria produce most of the energy required for cardiac function through oxidative phosphorylation. The electron transport chain (ETC) generates a proton motive force (Δp) across the mitochondrial inner membrane, comprised of an electrochemical potential gradient (Δψm) and a proton gradient (ΔpH), which is used to synthesize ATP. There are ions and metabolites, including Na^+^, inorganic phosphate (P_i)_, and pyruvate, that are co-transported across the inner mitochondrial membrane (IMM) with protons, necessitating regulation of proton gradients [1–4]. While ΔpH is important for energy transduction and ion transport, mitochondrial matrix pH (pH_mtx)_ in itself plays an important role in cell function. Changes in local pH can alter the protonation state of regulatory amino acid residues, thereby influencing enzyme conformation, catalytic activity, and substrate binding affinity. Additionally, calcium can complex with phosphate in the mitochondrial matrix. This buffering mechanism is highly pH-sensitive, with alterations in pH_mtx r_egulating mitochondrial free [Ca^2+^] [5, 6]. Despite its importance in energy and calcium homeostasis, pH_mtx r_emains difficult to assess in the intact heart.

Due to the oxidative nature of the myocardium, mitochondria are implicated in a range of cardiovascular disease states, including cardiac ischemia-reperfusion (I/R) injury. During ischemia, the cytosol becomes acidified due to the transition from aerobic to anaerobic metabolism resulting in lactate accumulation, as well as ATP hydrolysis, which contributes to an accumulation of protons [7, 8]. Upon reperfusion, cytosolic pH is rapidly normalized as the accumulated lactate is removed and ion transport processes are re-established [9]. Mitochondrial calcium overload and production of reactive oxygen species (ROS) during reperfusion putatively lead to induction of the mitochondria permeability transition and generalized energetic dysfunction, contributing to I/R injury [9, 10].

Understanding the magnitude and timing of changes in pH_mtx d_uring I/R will inform changes in oxidative phosphorylation and ion transport, as well as the production of ROS which is highly sensitive to the energetic state of the mitochondrion [11, 12]. Furthermore, drug delivery strategies are being developed to exploit the acidic microenvironment of ischemic tissue and the protonation properties of drugs, making pH an important consideration in targeted drug delivery [13]. Considering growing interest in mitochondrial-targeted therapeutics, it is important to characterize how pH_mtx c_hanges during I/R in order to extrapolate similar strategies to the mitochondrion.

Overall, time resolved determination of pH_mtx d_uring I/R will help establish how pH contributes to cardiomyocyte injury and cardioprotection, and inform the development of therapies. Although ^31^P nuclear magnetic resonance (NMR) has enabled quantification of cytosolic pH in models of I/R from the chemical shift of P_i [_14, 15], this approach requires long acquisition times and cannot resolve pH_mtx s_ince the intracellular P_i p_ool is predominantly cytosolic. Mitochondrial targeting of pH-sensitive probes like 2′,7′-biscarboxyethyl-5(6)-carboxyfluorescein (BCECF) and SNARF-1 in the intact heart is problematic because mitochondrial internalization requires transport against a large negative Δψm [16]. More recently, genetically encoded fluorescent sensors like SypHer targeted to the mitochondria enabled assessment of pH_mtx i_n cells and rodent brain [17–19]. Extending these approaches to measure pH_mtx i_n the intact heart would greatly improve our understanding of cardiac physiology and disease.

In addition to the therapeutic rationale of the present study, the recent expansion of fluorescence-based measurements of mitochondrial signaling mediators in the intact heart requires careful consideration of pH_mtx._ Studies using fluorescent mitochondrial-targeted Ca^2+^ [20–22] and ROS sensors [22, 23] have provided new insights into the role of mitochondria in cardiac I/R injury. However, most fluorescent probes are highly sensitive to pH requiring that these studies be performed under conditions where changes in pH are defined or minimized [20, 24]. Determination of pH would enable the interpretation of pH-sensitive fluorescent probes in models where pH changes are substantial such as I/R.

To address how mitochondrial matrix pH responds to I/R, we measured pH in the adult mouse heart during global no-flow I/R using a genetically encoded, mitochondrial-targeted pH probe SypHer3. Expanding upon prior studies from our lab [20, 24], here we also tested whether mitochondrial Ca^2+^ (Ca^2+^_m)_ rises during ischemia using the genetically encoded, mitochondrial-targeted Ca²⁺ sensor R-GECO1. Correcting R-GECO1 fluorescence for the effects of pH enabled the assessment of relative changes in Ca^2+^_m d_uring I/R, expanding the utility of fluorescence approaches for studying the intact heart and providing new insights into mitochondrial Ca^2+^ dynamics during I/R.

## Results

### Characterization of SypHer3 in isolated mitochondria

SypHer3 is a genetically encoded fluorescent sensor comprised of a circularly permuted green fluorescent protein (cpGFP) conjugated to a bacterial OxyR regulatory domain [17]. The OxyR regulatory domain has been mutated such that the probe is rendered redox-insensitive while retaining its pH sensitivity [17]. It has been employed to study pH gradients in zebrafish embryos [17], pH oscillations in active neurons [25], and pH during I/R in neurons and rodent brain [18]. SypHer3 has a p*K*_a o_f 7.8 and experiences a pH-dependent increase in fluorescence making it a strong candidate for studies of pH_mtx,_ particularly in I/R injury where pH changes are large [17]. To test this, SypHer3 was expressed to the mitochondrial matrix of cardiomyocytes in the adult mouse heart using adeno-associated virus (Fig. 1a). Its expression and mitochondrial localization was confirmed by confocal microscopy. In cardiomyocytes isolated from the adult mouse heart, green fluorescence due to SypHer3 co-localized with tetramethylrhodamine methyl ester (TMRM) confirming is expression and localization to the mitochondrial matrix (Fig. 1b). To assess the utility of SypHer3 for monitoring pH_mtx,_ mitochondria were isolated from the adult mouse heart and incubated at pH 7.8 without substrate and then titrated with HCl to decrease extramitochondrial pH which was recorded using a pH electrode while fluorescence was recorded using a fluorimeter (Fig. 1c). As pH decreased, SypHer3 fluorescence also decreased (Fig.1c-d) confirming its responsiveness to pH.

**Figure 1:**
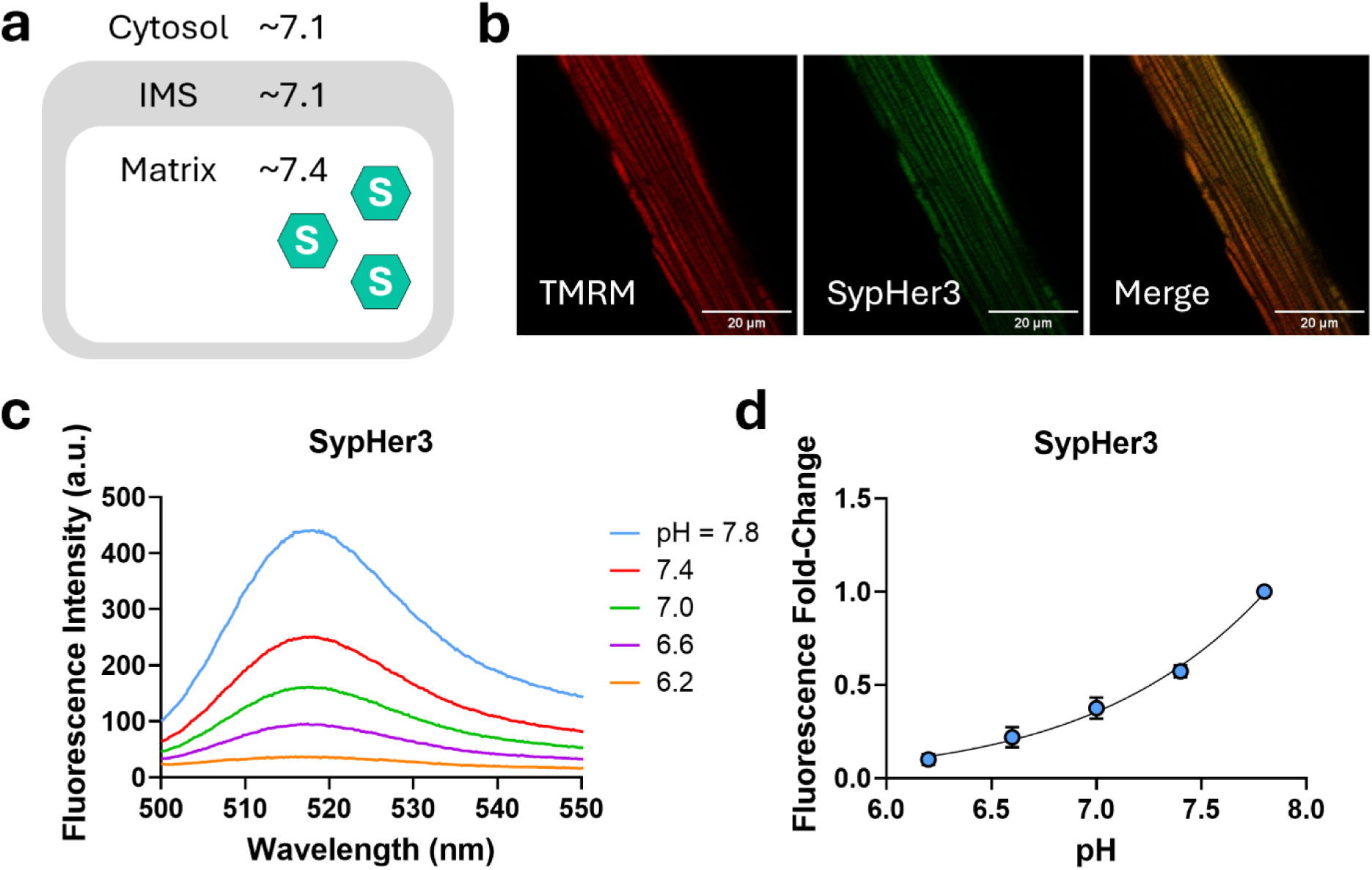
pH sensitivity of the genetically encoded pH sensor SypHer3. **a**, Schematic of a mitochondrion depicting SypHer3 localization (green hexagon). **b,** Isolated adult cardiomyocytes expressing SypHer3 incubated with 10 nM TMRM. **c,** Representative example of SypHer3 fluorescence in isolated heart mitochondria incubated without substrate (pH 7.8) and titrated with HCl to decrease extramitochondrial pH. **d,** The fold-change in SypHer3 fluorescence relative to pH 7.8 was calculated (n=4). Data are mean ± SD.

### Ǫuantitation of mitochondrial matrix pH from SypHer3 in vitro

To make a quantitative assessment of pH from SypHer3 fluorescence in the intact heart, SypHer3 fluorescence must first be calibrated against pH. To do this, a calibration was performed in isolated mitochondria to preserve the environment the probe is exposed to in vivo. Because the outer mitochondrial membrane (OMM) is highly permeable, the IMS is predicted to be in equilibrium with the extramitochondrial buffer [4]. However, even without added substrate, proton transport across the inner mitochondrial membrane (IMM) could sustain a pH gradient between the matrix and IMS. Since SypHer3 is localized to the mitochondrial matrix, a proton gradient across the IMM would preclude calibration of its fluorescence to pH based on the known extramitochondrial pH. Thus, isolated mitochondria were incubated with nigericin to eliminate ΔpH and equalize the pH of the matrix and the experimental buffer. As pH decreased from 7.8 to 6.2 by titration with HCl, SypHer3 fluorescence intensity (518 nm emission) declined in a non-linear fashion (Fig. 2a). To quantify pH_mtx f_rom SypHer3 fluorescence, SypHer3 fluorescence fold-change, rather than intensity, was calibrated against pH to normalize for differences in probe expression between biological replicates. This approach requires at least one experimental pH value to be known or assumed, against which the fluorescence fold-change can be calculated and calibrated. Here, SypHer3 fluorescence fold-change was calibrated relative to an assigned starting pH of 7.5 ± 0.06 (Fig. 2b) and the data fit with a continuous hinge function. This value was chosen because in the following experiments mitochondria were initially incubated in experimental buffer with pH ≈ 7.5. Mitochondria incubated in experimental buffer (pH 7.46 ± 0.03) were treated with nigericin to eliminate ΔpH and titrated with HCl to decrease the pH of the experimental buffer. The fold-change in fluorescence relative to the point after nigericin addition, where matrix pH was assumed to be in equilibrium with the experimental buffer (pH 7.46 ± 0.03), was determined (Fig. 2c). At each condition, pH was calculated from SypHer3 fold-change (Fig. 2d) using the calibration curve presented in Fig. 2b. SypHer3 fluorescence declined after the addition of nigericin consistent with a pH gradient across the IMM. To determine ΔpH, the difference between the calculated pH before nigericin and measured pH after nigericin was determined.

**Figure 2:**
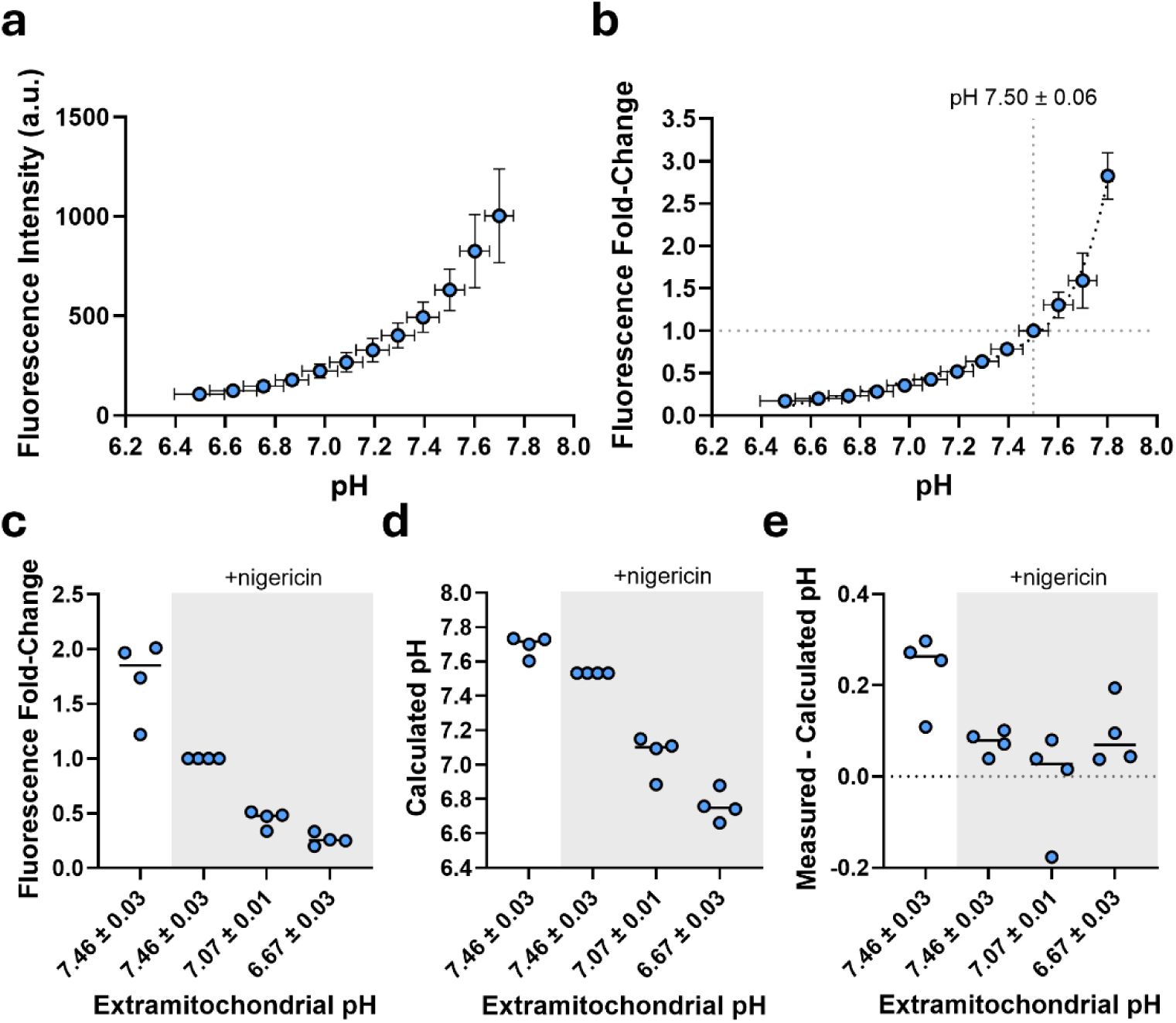
Calibration of SypHer3 fluorescence against matrix pH in isolated mitochondria. **a**, SypHer3 fluorescence intensity measured in isolated mitochondria incubated without substrate (pH 7.8) and with 1 μM nigericin, titrated with HCl to decrease extramitochondrial pH (n=3). **b,** The fold-change in SypHer3 fluorescence relative to pH 7.5 was calculated (n=3) and the data were fit with a continuous hinge function. **c,** Fold-change in SypHer3 fluorescence in isolated mitochondria incubated without substrate (pH 7.5), treated with 1 μM nigericin to dissipate ΔpH and titrated with HCl to decrease extramitochondrial pH (n=4). **d,** pH was calculated from fold-change by piecewise linear interpolation of the calibration curve shown in **b**. **e,** The difference in the measured and calculated pH was determined under each condition. Data are mean ± SD.

Assuming that the matrix pH was equal to the pH of the experimental buffer after nigericin (7.46 ± 0.03), the ΔpH where ΔpH = pH_mtx-_pH_IMS i_n non-respiring mitochondria incubated without substrate was 0.16 ± 0.06 units (n=4). The calculated pH after the addition of nigericin overestimated pH across conditions by < 0.1 units (Fig. 2e) demonstrating that nigericin effectively dissipated ΔpH and that this approach has utility for accurately estimating pH.

### Utility of SypHer3 in measuring mitochondrial matrix pH in the intact mouse heart

In order to determine whether SypHer3 has utility for measuring pH_mtx i_n the intact heart, we measured transmural SypHer3 fluorescence in the isolated perfused mouse heart. Transmural fluorescence was measured by positioning a light catheter in the left ventricle of the beating isolated perfused heart and exciting SypHer3 using a 488 nm fixed wavelength diode laser (Fig. 3a). Transmitted light was collected using an integrating sphere (Fig. 3b) connected to a rapid scanning spectrometer and fluorescence measured over the entire bandwidth of the spectrometer. Transmural fluorescence was recorded in hearts isolated from wild-type mice or mice expressing SypHer3 (Fig. 3c). Hearts expressing SypHer3 had fluorescence consistent with that observed in isolated mitochondria (Fig. 1c). Importantly, hearts from uninjected wild-type mice had negligible autofluorescence over the same bandwidth when assessed under the same conditions confirming that the fluorescence observed in hearts expressing SypHer3 was specific to SypHer3.

**Figure 3:**
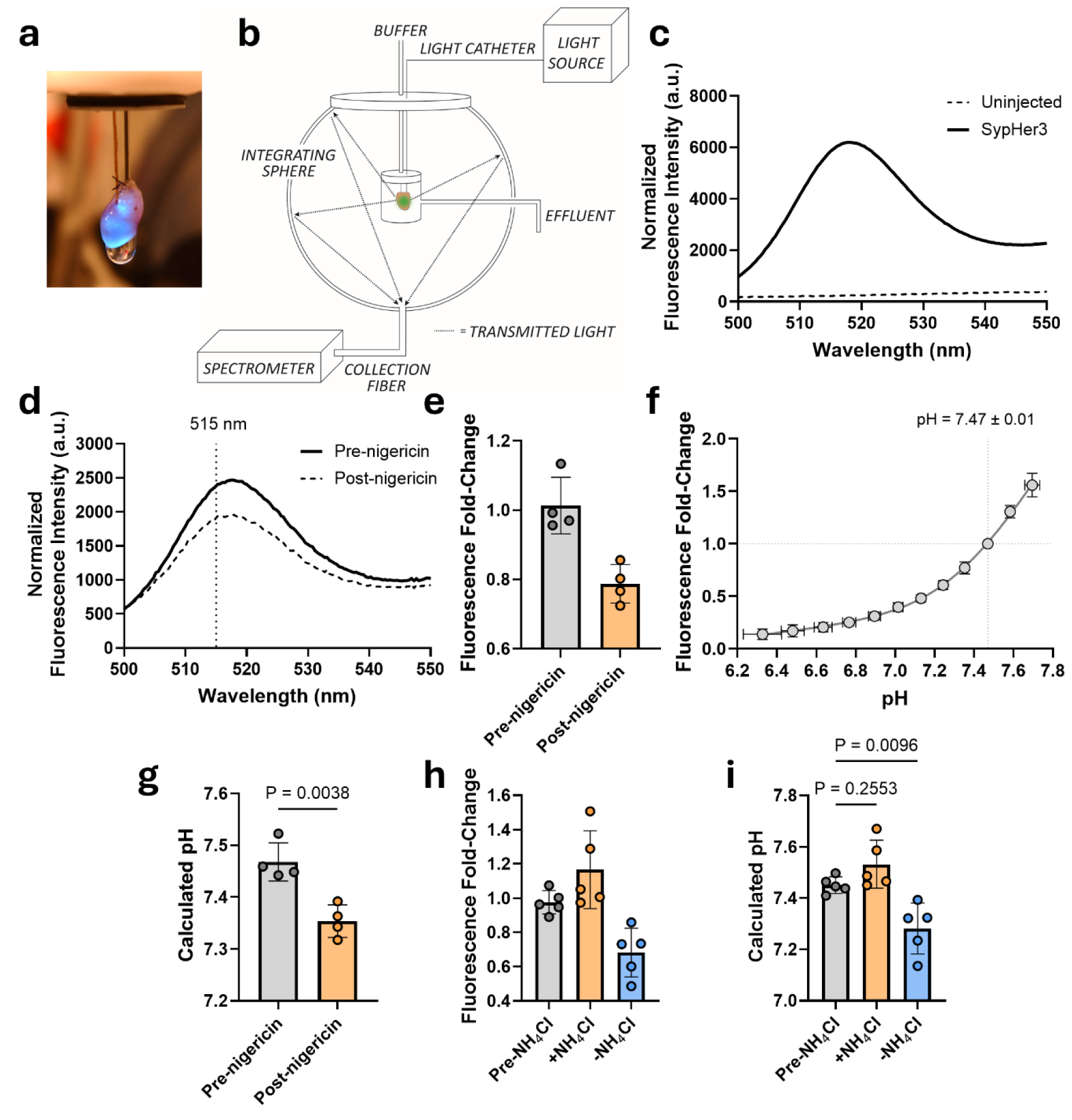
Utility of SypHer3 as a mitochondrial matrix pH sensor in the isolated perfused heart. **a**, Photograph of isolated perfused heart excited with a 488 nm laser. **b,** Schematic depicting the integrating sphere system used to measure transmural fluorescence of the isolated heart. **c,** Representative transmural fluorescence spectrum of an isolated heart from an uninjected wild-type mouse or wild-type mouse expressing SypHer3. **d,** Hearts were isolated from adult mice expressing SypHer3 and perfused by Langendorff perfusion. After 20 minutes, hearts were treated with 2 μM nigericin for 10 minutes. Representative SypHer3 fluorescence before and after nigericin is shown after correcting for primary inner-filter effects. **e,** The fold-change in fluorescence before nigericin and 10 minutes after nigericin was calculated relative to a control period (n=4). **f,** The calibration procedure described for mitochondria was repeated in the integrating sphere to mimic the optical conditions present in isolated perfused heart studies. Isolated mitochondria expressing SypHer3 were incubated without substrate and with 1 μM nigericin (pH 7.8) and titrated with HCl to decrease extramitochondrial pH. The fold-change in fluorescence was calculated relative to pH 7.47 ± 0.01 and the data fit with a continuous hinge function. **g,** pH was calculated from fold-change by piecewise linear interpolation of the calibration curve shown in **f**. A paired two-tailed t-test was performed to determine whether nigericin decreased pH. **h,** The isolated perfused heart was perfused for 20 minutes before perfusion with 10 mM NH_4C_l for 5 minutes followed by its washout. The fold-change in fluorescence relative to a control period was calculated (n=5). **i,** pH was calculated as described and a one-way ANOVA with Dunnett’s multiple comparisons used to test whether NH_4C_l pulse changed pH relative to control. Data are mean ± SD.

To determine whether SypHer3 responded as predicted to alterations in pH_mtx,_ hearts were treated with nigericin (2 µM) which facilitates K^+^/H^+^ exchange across the plasma membrane. The large extracellular K^+^ gradient is predicted to drive proton accumulation, resulting in cytosolic acidification. After making corrections for primary inner-filter effects as described in the Methods (supplemental Fig. 1a-c), the fold-change in fluorescence at 515 nm was determined pre-nigericin and after 10 minutes of treatment with nigericin, relative to a control period (Fig. 3e). There was a decrease in SypHer3 fluorescence with the addition of nigericin consistent with a decrease in pH due to cytosolic acidification resulting from plasmalemmal K^+^/H^+^ exchange. In order to quantify pH in this system, the calibration procedure described above was repeated in isolated mitochondria using the integrating sphere system to mimic the optical conditions during isolated heart studies (supplemental Fig. 1d). For quantitation in the intact heart, fold-change was calibrated to pH based on an assigned starting pH of 7.47 (Fig. 3f) and the data fit with a continuous hinge function. The starting pH_mtx w_as set to 7.47, given an experimental KH buffer with pH = 7.4 and an assumed cytosolic pH of 7.1-7.2 [26, 27] and ΔpH of 0.2-0.4 units in the isolated perfused heart [28, 29]. Based on this standard curve nigericin decreased pH_mtx b_y 0.11 ± 0.03 units (Fig. 3g).

To further test that SypHer3 responds as predicted to alterations in pH_mtx,_ hearts were subjected to an ammonium chloride (NH_4C_l) pulse to temporarily alkalinize the cytosol, followed by its washout to cause acidification [30, 31]. The fold-change in fluorescence pre-NH_4C_l, just after the addition of NH_4C_l, and 5 minutes after the washout of NH_4C_l were determined relative to a control period (Fig. 3h), and pH calculated from the standard curve presented in Fig. 3f. The addition of NH_4C_l had a modest impact on pH, whereas the washout of NH_4C_l led to a significant decrease in pH consistent with an acidification of the tissue (Fig. 3i). These results demonstrate that SypHer3 responds as predicted to changes in intracellular pH and has utility for measuring pH_mtx i_n the intact heart.

### Changes in mitochondrial matrix pH during cardiac ischemia-reperfusion injury

To understand how pH_mtx c_hanges during I/R, SypHer3 fluorescence was recorded in the intact heart subjected to global no-flow ischemia followed by 20 minutes of reperfusion and fluorescence recorded over time (Fig. 4a). SypHer3 fluorescence was measured at discrete time points and corrected for primary inner-filter effects (Fig. 4b). The fold-change in fluorescence was calculated relative to the time just before ischemia, and is shown without and with correction for primary inner-filter effects (Fig. 4c). The fold-change in SypHer3 relative to control was determined after correction for primary inner-filter effects and used to calculate pH (Fig. 4d) from the calibration curve presented in Fig. 3f using an assigned starting matrix pH = 7.47. SypHer3 fluorescence declined during ischemia and recovered to control over the first 5 minutes of reperfusion, consistent with a decline in pH, followed by a rapid recovery.

**Figure 4:**
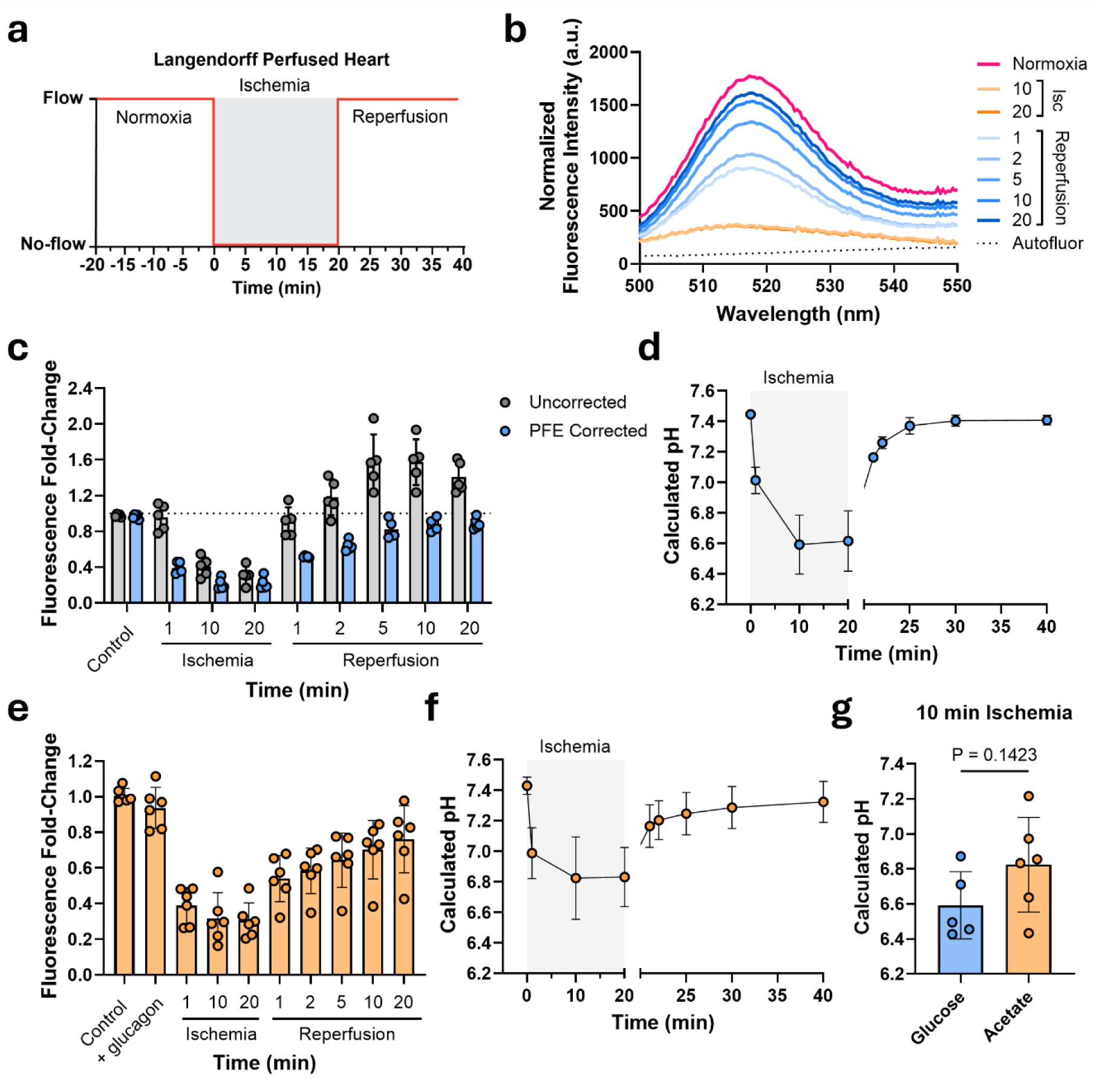
Changes in mitochondrial pH during global no-flow ischemia-reperfusion. **a**, Schematic of the Langendorff perfusion protocol. **b,** The isolated heart expressing SypHer3 was perfused with glucose-containing buffer and subjected to 20 minutes of ischemia followed by 20 minutes of reperfusion. SypHer3 fluorescence spectra are shown at discrete time points after correction for primary inner-filter effects (representative). Autofluorescence was measured in an uninjected wild-type heart is shown. **c,** The fold-change in SypHer3 fluorescence relative to the normoxic control was calculated prior to optical correction, and after correction for primary inner-filter effects (PFE) (n=5). **d,** pH was calculated from **c** by piecewise linear interpolation of the calibration curve shown in Fig. 3f. **e,** The isolated heart expressing SypHer3 was perfused without glucose and administered 2 ug/mL glucagon three minutes prior to 20 minutes of ischemia to glycogen deplete the heart. The fold-change in fluorescence was calculated relative to the normoxic control (n=6). **f,** pH was calculated from **e** as described above (n=6). **g,** pH after 10 minutes of ischemia in hearts perfused with glucose (n=5) or acetate (n=6). An unpaired two-tailed t-test was used to test whether pH was statistically different. Data are mean ± SD.

Next we tested whether glycogen depletion impacted pH_mtx d_uring I/R. During ischemia, anaerobic glycolysis becomes the primary source of cellular ATP using glucose released from glycogen stores, contributing to intracellular acidosis [9]. By depleting glycogen prior to ischemia, less glucose is available for glycolysis resulting in a smaller decrease in cytosolic pH [20, 32]. Thus, we predicted that the pH_mtx w_ould decrease less during ischemia following glycogen depletion. Hearts were perfused with buffer containing 5 mM sodium acetate substituted for glucose and administered 2 µg/mL glucagon three minutes prior to 20 minutes of global no-flow ischemia, followed by reperfusion. SypHer3 fluorescence was recorded at discrete time points and fold-change calculated after correction for primary inner-filter effects (Fig. 4e), and pH calculated as above (Fig. 4f).

Although the pH_mtx a_fter 10 minutes of ischemia trended higher in hearts perfused with acetate-containing buffer compared to those perfused with glucose-containing buffer (Fig. 4g), pH_mtx w_as not significantly different between the two substrate conditions. Here, we demonstrate that pH_mtx c_hanges dynamically during I/R injury, declining during ischemia and rapidly recovering upon reperfusion.

### Changes in mitochondrial calcium during cardiac ischemia-reperfusion injury

Our lab has recently expanded fluorescence based measurements of calcium (Ca^2+^) in the isolated perfused heart. Using the genetically encoded mitochondrial-targeted Ca^2+^ sensor R-GECO1 we investigated changes in mitochondrial Ca^2+^ (Ca^2+^_m)_ during cardiac I/R injury [20, 24, 33]. However, as with most fluorescent probes, R-GECO1 is sensitive to pH requiring these studies to be performed under conditions where changes in intracellular pH were attenuated by glycogen depletion. The ability to measure pH during I/R using SypHer3 will enable the interpretation of changes in R-GECO1 fluorescence without altering substrate conditions, better mimicking the in vivo condition. To assess how Ca^2+^_m c_hanges during I/R in the presence of glucose and endogenous glycogen, hearts expressing R-GECO1 were perfused with glucose-containing KH buffer and subjected to ischemia and reperfusion and fluorescence recorded at discrete time points. Fluorescence intensity was measured at 650 nm and corrected for primary inner-filter effects (Fig. 5a) as previously described [20]. In the presence of glucose, R-GECO1 fluorescence decreased early in ischemia (Fig. 5a), consistent with a similar study from our lab [20]. However, as shown in Fig. 4d, pH_mtx d_eclines significantly over this time period. Zhao et al. previously characterized the relationship between R-GECO1 fluorescence and pH in the presence or absence of Ca^2+^ using purified protein [34]. The authors found that R-GECO1 fluorescence declined as pH decreased from 7.6 to 6.0 in both Ca^2+^-bound or Ca^2+^-free states (Fig. 5b) [34]. Thus, a decrease in pH_mtx w_ould decrease R-GECO1 fluorescence irrespective of Ca^2+^ binding state. To account for the effect of pH on R-GECO1 fluorescence, pH_mtx w_as determined at each time point from SypHer3 fluorescence fold-change as described above (Fig. 4d, Table 1). A pH-dependent correction factor f(pH_x)_ was then derived (Fig. 5c) from the calibration of R-GECO1 fluorescence against pH (Fig. 5b) for each experimental pH value using data adapted from Zhao et al., as described in the *Methods* (Fig. 5c) [34]. R-GECO1 fluorescence was then divided by the correction factor corresponding to the measured pH_mtx,_ giving a corrected fluorescence intensity (Fig. 5a, Table 1). The fold-change in corrected fluorescence was determined relative to control (Table 1). R-GECO1 fluorescence was significantly higher than control after 20 minutes of ischemia, returning to control levels within the first minute of reperfusion (Fig. 5a, Table 1).

**Figure 5:**
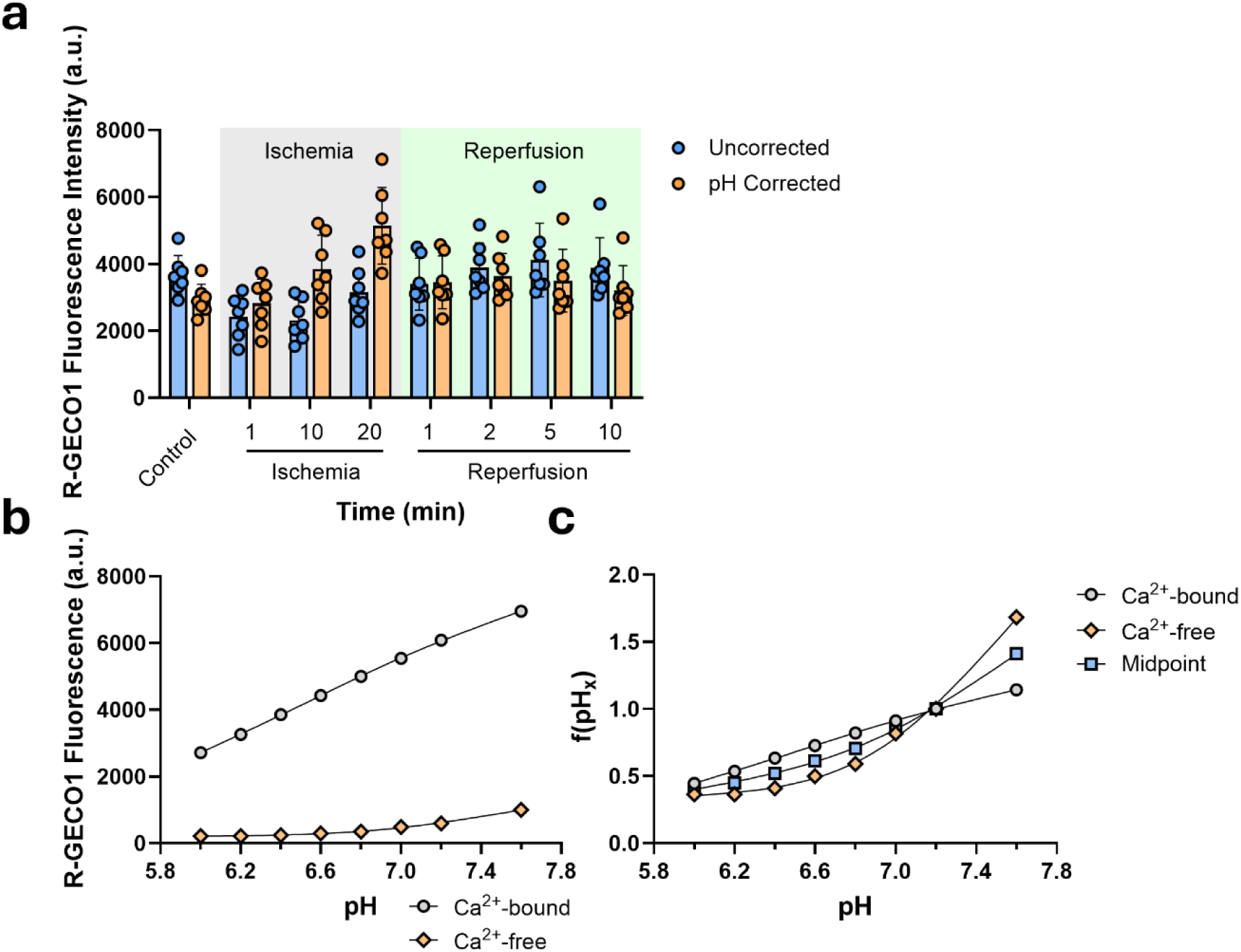
Effect of pH on R-GECO1 fluorescence during global no-flow ischemia-reperfusion. **a**, The isolated heart expressing R-GECO1 was perfused with glucose-containing buffer and subjected to 20 minutes of ischemia followed by 20 minutes of reperfusion. R-GECO1 fluorescence was measured over time and corrected for primary inner-filter effects. R-GECO1 fluorescence is shown before (uncorrected) and after correction for pH as described in text. Data are mean ± SD. **b,** R-GECO1 fluorescence was measured in purified protein as a function of pH with (Ca^2+^-bound) or without (Ca^2+^-free) 10 mM CaCl_2._ Data adapted from Zhao et. al. (2011). **c,** A pH-dependent correction factor f(pH_x)_ was determined by taking the ratio of R-GECO1 fluorescence at a given pH relative to pH 7.2 for both the Ca²⁺-bound and Ca²⁺-free states (derived from **b**), and the resulting data fit with a smoothing spline curve. The midpoint was then calculated as the mean f(pH_x)_ of the Ca^2+^-bound and Ca^2+^-free functions and fit with a smoothing spline curve.

**Table 1:** Correction of R-GECO1 fluorescence for changes in mitochondrial pH during ischemia-reperfusion. Changes in pH_mtx (_n=5) calculated from SypHer3 fluorescence fold-change and R-GECO1 fluorescence (n=7) in the isolated perfused mouse heart during global ischemia-reperfusion. Fluorescence intensity, F(pH_x)_, and corrected fluorescence intensity (F_corr)_ were determined as described in text. Fold-change relative to control was calculated using F_corr._ A one-way ANOVA with Dunnett’s multiple-comparisons was used to compare pH-corrected R-GECO1 fluorescence (F_corr)_ to control. An * represents statistical significance (P > 0.05). Data are mean ± SD.

|  | **Time (min)** | **pH** | **f(pH<sub>x</sub>)** | **R-GECO1 F(pH<sub>x</sub>) (a.u.)** | **R-GECO1 F<sub>corr</sub> (a.u.)** | **Fold-Change** | **P value** |
|---|---|---|---|---|---|---|---|
|  | Control | 7.45 ± 0.01 | 1.25 | 3676 ± 582 | 2934 ± 464 | - | - |
| Ischemia | 1 | 7.01 ± 0.09 | 0.86 | 2428 ± 623 | 2826 ± 725<sup>ns</sup> | 0.97 ± 0.21 | 0.996 |
|  | 10 | 6.59 ± 0.19 | 0.60 | 2325 ± 607 | 3858 ± 1006<sup>ns</sup> | 1.35 ± 0.42 | 0.305 |
|  | 20 | 6.62 ± 0.20 | 0.61 | 3155 ± 703 | 5143 ± 1145* | 1.77 ± 0.39 | 0.008 |
| Reperfusion | 1 | 7.16 ± 0.01 | 0.98 | 3398 ± 777 | 3453 ± 790<sup>ns</sup> | 1.19 ± 0.28 | 0.471 |
|  | 2 | 7.26 ± 0.04 | 1.07 | 3898 ± 729 | 3638 ± 681<sup>ns</sup> | 1.26 ± 0.27 | 0.251 |
|  | 5 | 7.37 ± 0.05 | 1.18 | 4122 ± 1100 | 3499 ± 934<sup>ns</sup> | 1.22 ± 0.35 | 0.626 |
|  | 10 | 7.40 ± 0.04 | 1.21 | 3884 ± 899 | 3207 ± 742<sup>ns</sup> | 1.11 ± 0.26 | 0.888 |

## Discussion

Mitochondrial matrix pH (pH_mtx)_ is central to cardiac energetics and has implications for enzyme activity, energetic status, cellular redox balance, metabolite flux, and ion transport. Until now it has not been possible to quantify pH_mtx i_n the intact heart due to challenges targeting pH probes to the mitochondrion [16]. Here, we used the genetically encoded, mitochondrial matrix targeted pH sensor SypHer3 to measure pH_mtx i_n the isolated mouse heart. This approach offers the advantage of stable expression and localization, overcoming uncertainties related to the uptake and compartmentalization of mitochondrial targeted dyes and probes.

The primary objective of this study was to understand how pH_mtx c_hanges during I/R. The isolated heart was subjected to global no-flow I/R and pH_mtx e_stimated from SypHer3 fold-change relative to the normoxic control over time by extrapolating the calibration of SypHer3 fluorescence fold-change against pH established in isolated mitochondria. The assessment of fold-change rather than absolute intensity provides normalization for probe expression between biological replicates, but requires the starting pH value to be known or assumed. Although absolute fluorescence intensity could, in principle, provide accurate pH quantitation, this would require knowledge of the precise concentration of SypHer3 sampled using our ex vivo optical system which is unknown and difficult to determine due to the complex optical geometry of the heart. Ǫuantitation herein assigned a starting pH_mtx o_f 7.47 based on an assumed cytosolic pH between 7.1-7.2 [26, 35, 36] and ΔpH between 0.2-0.4 units in heart mitochondria [4, 19, 28]. However, reported estimates of cytosolic pH and ΔpH vary depending on experimental conditions, introducing uncertainty into the absolute pH values estimated using this approach. To facilitate application of this method to other experimental models, SypHer3 fold-change can be calibrated against a range of starting pH values. As examples, fold-change calibrations were generated using starting pH values of 7.35 ± 0.01, 7.47 ± 0.01, and 7.58 ± 0.02 (supplemental Fig. 2a). As the predicted starting pH increased from 7.35 to 7.58, the calculated pH values increased, however the trends observed were unaffected with pH declining during ischemia and recovering to control values over the first 5 minutes of reperfusion (supplemental Fig. 2b). Assuming a starting pH of 7.47, pH_mtx d_ecreased by between 0.83 ± 0.19 units (n=5) after 20 minutes of ischemia and recovered to 7.16 ± 0.01 (n=5) within the first minute of reperfusion, with pH_mtx c_ontinuing to recover over the duration of reperfusion.

The magnitude and temporal behavior of pH_mtx u_pon reperfusion was previously unknown and its recovery has implications for ATP production, reactive oxygen species (ROS) production, ion and metabolite transport, calcium phosphate buffering, and the mitochondria permeability transition. The decline in pH_mtx d_uring ischemia can be explained in part by the inhibition of respiration due to oxygen depletion and equilibration with the IMS, and acidified cytosol which declines from ∼7.1 at normoxia to ∼6.2 during ischemia [35, 36]. Upon reperfusion, there was a rapid recovery of pH_mtx c_orresponding to the recovery of cytosolic pH revealed by ^31^P NMR studies of the isolated heart [26, 32, 35], consistent with a recovery of ΔpH. A rapid recovery of ΔpH, along with the re-establishment of mitochondrial membrane potential (Δψm) early in reperfusion [37] supports re-energization, as well as re-establishment of proton-coupled ion transport processes (e.g., Na^+^, K^+^, and P_i)_ [1–3].

This approach enabled us to address the impact of pH on R-GECO1, a mitochondrial Ca^2+^ sensor. Prior studies with R-GECO1 required that the heart be depleted of glycogen to minimize intracellular acidosis associated with glycolysis during ischemia in order to isolate changes in fluorescence due to Ca^2+^ [20]. These studies saw an increase in Ca^2+^_m d_uring ischemia, mediated in part by a Ca^2+^ uptake pathway distinct from the voltage-dependent mitochondrial calcium uniporter (MCU) [20]. While glycogen depletion has been shown to attenuate the rise in lactate [32] and cytosolic acidosis [20], we found that it did not fully prevent the decline in pH_mtx (_Fig. 4g). Thus, in order to recapitulate the finding that Ca^2+^_m r_ises during ischemia without altering substrate conditions, we sought to correct R-GECO1 fluorescence for changes in pH_mtx._ By measuring pH_mtx a_nd R-GECO1 fluorescence during I/R in paired studies, and extrapolating an existing calibration of R-GECO1 fluorescence against pH [34], we were able to correct R-GECO1 fluorescence for pH to assess relative changes in Ca^2+^_m._ Consistent with prior studies from our lab, Ca^2+^_m r_ises during ischemia and declines to control levels upon reperfusion [20]. It is important to acknowledge that the relationship R-GECO1 fluorescence and pH depends on the Ca^2+^ occupancy of R-GECO1. Because occupancy could not be determined experimentally, we used the arithmetic mean of the fully Ca²⁺-bound and Ca²⁺-free correction factors derived from data collected by Zhao et. al. [34], which define the upper and lower bounds of the pH correction. As f(pH_x)_ varied only modestly over the observed pH_mtx r_ange (6.5–7.5) (Fig. 5c), uncertainty due to occupancy is expected to have a modest impact on the corrected signal. Although this approach was used to address the impact of pH on RGECO1, this could be extrapolated to any pH-sensitive probe. Indeed, without correcting for pH, Ca^2+^_m a_pparently declines early in ischemia, providing rationale for this work. The NEMO family of fluorescent sensors is a new class of Ca^2+^ probes with a range of affinities and fluorescent properties, and a lower sensitivity to pH, that could be considered in future studies of Ca^2+^ in the intact heart [38].

Under typical conditions, mitochondrial Ca²⁺ can be complexed or precipitated with phosphate or polyphosphate rendering it osmotically inactive [5, 6]. It has been shown that Ca^2+^ phosphate is less soluble as pH declines [5] implying that during ischemia when pH_mtx i_s low Ca^2+^ phosphate could dissociate, contributing to the increase in Ca^2+^_m o_bserved. Indeed, considering that Δψm falls below-100 mV during ischemia [37], potentially impacting MCU activity [39], and that knockout of MCU does not block the rise in Ca^2+^_m,_ it is possible that the increase in Ca^2+^_m c_ould be explained in part by dissociation of Ca^2+^ phosphate. This possibility is also consistent with the rapid decline in Ca^2+^_m u_pon reperfusion, which may be explained by rapid re-sequestration as pH returns to normoxic levels (Fig. 4d). Future studies will investigate the impact of pH on mitochondrial Ca^2+^ uptake and phosphate-linked Ca^2+^ buffering.

In summary, we developed a fluorescence approach for measuring pH_mtx i_n the isolated perfused moused heart providing estimates of pH_mtx a_nd new insights into Ca^2+^ dynamics during I/R. This approach represents a major advancement in fluorescence-based studies of the intact heart expanding our toolkit for studying mitochondrial mechanisms of cardiac physiology and disease and providing novel insights into cardiac I/R injury.

**Supplemental Figure 1:**
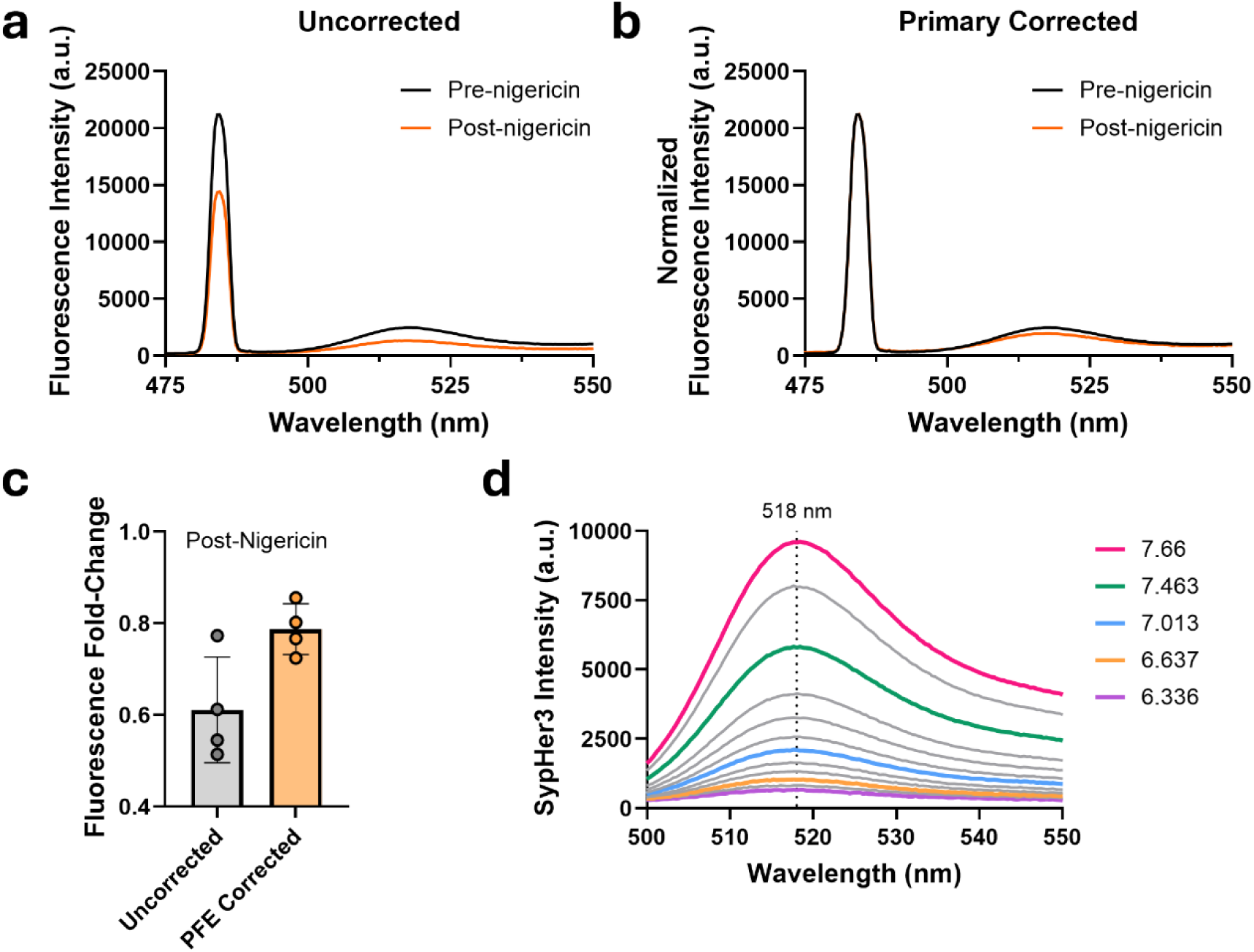
Measuring SypHer3 fluorescence and pH in the isolated perfused mouse heart. **a**, Hearts were isolated from adult mice expressing SypHer3 and perfused by Langendorff perfusion. After 20 minutes, hearts were treated with 2 μM nigericin for 10 minutes. SypHer3 fluorescence was measured before and after nigericin (representative). **b,** SypHer3 fluorescence after normalizing light transmission at 488 nm to that of the control condition to correct for primary inner-filter effects (representative). **c,** The fold-change in SypHer3 fluorescence after nigericin relative to control was determined from the raw fluorescence intensity, and from fluorescence intensity after primary filter correction (PFE) (n=4). **d,** Isolated mitochondria expressing SypHer3 were incubated in the integrating sphere without substrate and with 1 μM nigericin (pH 7.8) and titrated with HCl to decrease extramitochondrial pH. SypHer3 fluorescence was measured at each pH value (representative).

**Supplemental Figure 2:**
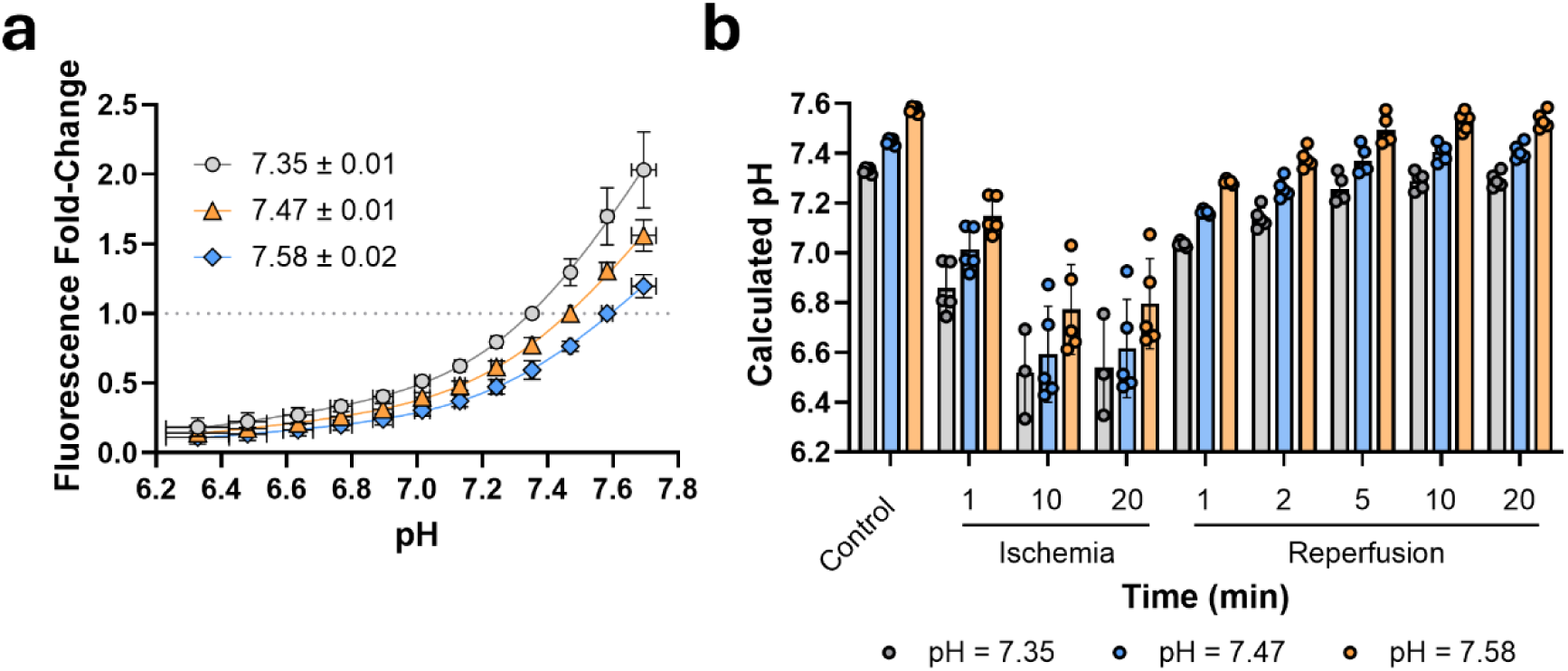
Ǫuantitation of mitochondrial matrix pH during global no-flow ischemia-reperfusion. **a**, Isolated mitochondria expressing SypHer3 were incubated in the integrating sphere without substrate and with 1 μM nigericin (pH 7.8) and titrated with HCl to decrease extramitochondrial pH. The average fold-change in SypHer3 fluorescence was calculated relative to pH 7.35, 7.47, and 7.58 and the data fit with a continuous hinge function (n=3). **b,** The isolated heart expressing SypHer3 was perfused with glucose-containing buffer and subjected to 20 minutes of ischemia followed by 20 minutes of reperfusion. The fold-change in SypHer3 fluorescence relative to normoxic control was calculated after correction for primary inner-filter effects and pH calculated by piecewise linear interpolation using the corresponding calibration of SypHer3 relative to pH shown in **a** (n=5). Data are mean ± SD.

## Methods

### Ethical approval and use of animals

All animal protocols were approved by the Animal Care and Use Committee National Heart, Lung, and Blood Institute, National Institutes of Health (Bethesda, MD) and were performed in accordance with guidelines established by the Animal Welfare Act. Male and female C57Bl/6N mice were bred in-house and housed separately with ad libitum access to food and water.

### Expression of SypHer3 by adeno-associated virus

AAV9-CMV-mito-SypHer3 and AAV9-CMV-mito-R-GECO1 were obtained from Vector Biolabs and stored at –80°C. Neonatal mice postnatal day 3-5 received intrathoracic injections of 35 μL of AAV as previously described [20]. AAV was diluted in sterile saline for a final concentration of ∼5 x 10^11^ genome copies (GC) per injection. Neonatal pups were briefly cryo-anesthetized before injection and animals identified by toe clip. Animals were placed on a heating pad at 37°C for monitoring until mobility was regained and then returned to their parents.

### Primary adult cardiomyocyte isolation

Cardiomyocytes were isolated from isolated perfused mouse hearts by constant pressure perfusion as previously described [40]. Isolated hearts were perfused with KH buffer for 20 minutes and then transitioned from standard KH buffer to a relaxation buffer containing (in mM) 120 NaCl, 15 NaHCO_3,_ 10 KHCO_3,_ 10 glucose, 1.2 KH_2P_O_4,_ 4.7 KCl, 1.2 MgSO_4,_ 30 taurine, 12.5 BDM, and 0.4 EGTA. Hearts were then digested for 10-15 minutes with digestion buffer containing (in mM) 120 NaCl, 15 NaHCO_3,_ 10 KHCO_3,_ 10 D-glucose, 1.2 KH_2P_O_4,_ 4.7 KCl, 1.2 MgSO_4,_ 30 taurine, 12.5 BDM, 0.2 mg mL^-1^ Liberase (5401119001, Roche), and 12.5 µM CaCl_2 a_t a flow rate of 2-3 mL min^-1^. At the end of the digestion period, the heart was removed from the perfusion system and digestion stopped with a stop buffer containing (in mM) 120 NaCl, 15 NaHCO_3,_ 10 KHCO_3,_ 10 glucose, 1.2 KH_2P_O_4,_ 4.7 KCl, 1.2 MgSO_4,_ 30 taurine, 12.5 BDM, and 1 mg mL-1 fatty acid free BSA. The heart was teased apart gently with forceps. for 5-8 minutes. Cells and debris were transferred to a 15 mL falcon tube and gravity pelleted for 2 minutes. The supernatant containing cardiomyocytes was transferred to a new 15 mL falcon tube and gravity pelleted for 10 minutes. The supernatant was removed and discarded and replaced with 5 mL Stop Buffer supplemented with 1 mM CaCl_2._ Cells were resuspended gently using a Pasteur pipette with the tip removed. Cells were allowed to gravity pellet again for 10 minutes and then resuspended in experimental buffer containing (in mM) 120 NaCl, 15 NaHCO_3,_ 10 KHCO_3,_ 10 glucose, 1.2 KH_2P_O_4,_ 4.7 KCl, 1.2 MgSO_4,_ 30 taurine, 12.5 BDM, and 1 CaCl_2._

### Confocal microscopy

Cardiomyocytes cells were plated on laminin-coated 35 mm glass bottom dishes (P35G-0-20-C, MatTek Corporation). Cells were incubated with or without 10 nM TMRM for 20 min at 37°C and then washed and refreshed with experimental buffer. Cells were imaged on Leica DMI6000B-CS using a 63×1.32 oil objective. TMRM was excited at 550 nm (7% power) and emission collected between 560-700 nm. SypHer3 was excited at 488 nm (0.8% power) and emission collected between 495-545 nm emission. Images were collected using Leica Imaging Software and processed using Fiji in ImageJ.

### Isolation of mitochondria from mouse heart

Mitochondria were isolated by mechanical homogenization and trypsin digestion and separated by differential centrifugation in isolation buffer containing (in mM) 225 mannitol, 75 sucrose, 5 MOPS, 0.5 EGTA and 2 taurine (pH 7.25) as previously described [41]. The final pellet was resuspended in 400 µL of isolation buffer and mitochondrial protein measured by Bradford assay. Mitochondria were incubated in 1.5 mL buffer containing (in mM) 120 KCl, 10 Tris-HCl, 5 MOPS, 5 KH_2P_O_4._ The pH of the buffer was adjusted with HCl or KOH as indicated in the text. A micro pH probe connected to a pH meter (Sartorius) was used to measure pH.

### Langendorff perfusion of the isolated mouse heart

Male or female C57BL/6N mice between 10 and 16 weeks of age were anesthetized with sodium pentobarbital (50 mg) and administered 100 units of USP-grade heparin by intraperitoneal injection. After determining proper depth of anesthesia, the heart was excised and submerged in ice-cold KH buffer containing (in mM) 120 NaCl, 25 NaHCO3, 4.7 KCl, 1.75 CaCl_2,_ 1.2 MgSO_4,_ 1.2 KH_2P_O_4 a_nd 10 d-glucose filtered before use. After excision, the heart was cannulated and retrogradely perfused at constant pressure (90 mm Hg) with KH buffer maintained at 37°C and bubbled with 95% O_2/_5% CO_2 (_pH 7.4). The left atrium was removed and a 220 µm light catheter (FIP200220240, Molex) positioned inside the left ventricular cavity. The heart was secured in a water-jacketed glass chamber maintained at 37°C. The effluent flow rate was measured continuously using a flow-through sensor (1PXN, Transonic) positioned before the heart, coupled to a Tubing Flow Module (TS410, Transonic). Temperature was measured using a flexible temperature microprobe (IT-18, AD Instruments). Flow and temperature signals were digitized via a PowerLab interface (AD Instruments). Flow was recorded at 1,000 samples per second, and heart rate calculated based on the sinusoidal flow rate with the peak of each wave taken to be one beat and the number of peaks calculated over time.

### Isolated perfused heart studies

Nigericin was prepared as a 10 mM stock solution in DMSO and diluted in KH before being infused at 1% of the coronary flow rate to achieve the desired final concentration (2 μM).

The ammonium chloride (NH_4C_l) pulse was achieved by perfusing hearts normally for 20 minutes before perfusion with KH buffer containing 10 mM NH_4C_l. Hearts were perfused with NH_4C_l for 5 minutes before transition back to standard KH buffer for up to 5 minutes. Fluorescence was determined just before the transition to NH_4,_ after 5 minutes of NH_4,_ and 1 minute after the washout was initiated.

To deplete the isolated perfused heart of glycogen, glucose was replaced with 5 mM sodium acetate. Three minutes before global no-flow ischemia, glucagon (G2044, Sigma-Aldrich) was infused at 10% of the coronary flow rate for a final concentration of 2 μg ml^−1^ as previously described [20].

### Transmural optical spectroscopy

A 488 nm fixed wavelength diode laser (Cobalt 06-MLD, Hübner Photonics) was used to excite SypHer3. A 535 nm laser (Ocean Optics) was used to excite R-GECO1. To transmit light, the light catheter was connected to both the laser and a 560-nm broadband white light source (LCS 0560-68-22, Mightex) using a bifurcated fiber bundle (BFY1000HS02, Thor Labs). Transmitted light was collected using a multimode optical fiber (FT1000UMT, Thor Labs) fitted to the bottom port of an integrating sphere (LabSphere) and measured by a cooled, rapid-scanning spectrometer (VIS-A-S-100, Wasatch Photonics). A variable attenuator (FOFMS, Thorlabs) with a 515 nm long pass filter (FGL515M, Thorlabs) was incorporated between the integrating sphere and spectrometer to selectively attenuate the transmitted laser light enabling monitoring of the excitation wavelength without saturating the detector. Light intensity was recorded at 1,024 points between 400 and 800 nm at one sample per second using a custom LabVIEW-based program [42]. The incident light intensity (I_0)_ and dark current of the spectrometer were collected at the start of each day.

### Spectral analysis

Fluorescence was measured in isolated mitochondria using a spectrofluorometer (RF-6000, Shimadzu) and recording using LabSolutions RF. Two consecutive fluorescence spectra were averaged under each experimental condition and peak intensity measured at 515 nm. For isolated heart studies, fluorescence was measured using rapid scanning spectrometer (VIS-A-S-100, Wasatch Photonics) coupled to an integrating sphere (LabSphere) and recorded using a custom LabView-based program [42]. Between 15-30 fluorescence spectra were averaged prior to analyses. For the nigericin, NH_4C_l pre-pulse, and I/R studies, two consecutive control periods (at least 15 spectra each) were analyzed to establish a baseline control fluorescence intensity, which was used to calculate the experimental fold-change in fluorescence.

For isolated heart studies, excitation intensity was measured at 484 nm, and SypHer3 fluorescence emission recorded at 515 nm. Two major considerations in transmural fluorescence spectroscopy are the impact of primary and secondary inner-filter effects. When the isolated heart was administered nigericin, light transmission between 475-575 nm changed (supplemental Fig. 1a) consistent with primary inner-filter effects. Primary inner-filter effects arise when the tissue chromophores absorb the excitation light before it reaches the fluorophore, reducing the effective excitation intensity and impacting fluorescence emission independent of biochemical changes [43]. Fluorescence emission is proportional to fluorescence excitation [43], thus, the excitation intensity (484 nm) under the control condition (Ex_1_) and at each experimental condition (Ex_x_) was determined, and acquired fluorescence emission (515 nm) (Em_x_) multiplied by the ratio of Ex_1_/Ex_x_ to yield the corrected emission (Em_P_) (**Eq. 1**). This procedure normalizes excitation intensity to the control condition and scales the measured fluorescence emission to compensate for differences in excitation intensity (supplemental Fig. 1b) [20, 21, 43].

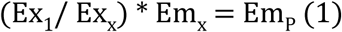

In this study, we did not attempt to correct for secondary inner-filter effects, which arise when tissue chromophores absorb fluorophore emission. Although tissue absorption is expected to substantially influence absolute fluorescence intensity, our analysis focused on changes in fluorescence intensity between experimental conditions. Transmission at 515 nm relative to control decreased during ischemia and increased upon reperfusion.

These changes likely reflect a combination of altered optical path length resulting from changes in tissue geometry (including myocardial thickening and edema) and changes in tissue absorbance associated with myoglobin and mitochondrial cytochromes [44, 45]. However, white-light transmission measurements reflect light traversing the entire heart wall, whereas fluorescence emission originates within the myocardium and is subject to an optical path that depends on the depth and distribution of the fluorescent signal.

Consequently, changes in bulk tissue transmission cannot be directly translated into changes in fluorescence emission attenuation, and applying a quantitative secondary inner-filter correction based on these measurements would require assumptions about photon path length that cannot be readily established in this preparation. To address this limitation, we measured SypHer3 fluorescence at 515 nm, a wavelength that is blue-shifted relative to its emission maximum (518 nm) where absorption by myoglobin and the spectral shoulder of cytochrome *c* is lower relative to 518 nm [44].

### Correction of R-GECO1 Fluorescence for pH-Dependent Effects

Because R-GECO1 fluorescence is pH-sensitive, fluorescence signals were corrected for pH-dependent effects using an empirical normalization factor derived from published calibrations of R-GECO1 fluorescence as a function of pH. The relationship between R-GECO1 fluorescence and pH for both the Ca²⁺-bound and Ca²⁺-free states was extrapolated from measurements of R-GECO1 fluorescence as a function of pH from Zhao et al. (Fig. 5b) [34]. For each state, a pH-dependent correction factor, f(pH_x)_, was calculated as the ratio of R-GECO1 fluorescence at a given pH relative to that at pH 7.2 (Fig. 5c). If R-GECO1 were exclusively bound or free, corrected fluorescence intensity (Fcorr) could be calculated by dividing the measured fluorescence at a given pH (Fm(pHx)) by f(pHx) using **Eq. 2**.

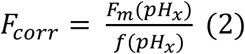

However, though the pH dependence of fluorescence is similar for the Ca²⁺-bound and Ca²⁺-free states over a physiological pH range (pH 6.4–7.4), the two relationships are not identical (Fig. 5b-c) [34]. Thus, the effect of pH on R-GECO1 fluorescence depends on the Ca^2+^ occupancy, which could not be determined under the present experimental conditions. To estimate the contribution of pH to the measured fluorescence signal, an intermediate correction function was calculated as the mean of the correction factors derived from the Ca²⁺-bound and Ca²⁺-free states at each pH value (Fig. 5c). The midpoint correction factor corresponding to an experimental pH value was then used to correct R-GECO1 fluorescence according to **Eq. 2**.

Importantly, the Ca²⁺-bound and Ca²⁺-free calibration curves define the upper and lower bounds of f(pH_x)_ and the true correction must fall between these limits. By averaging the f(pH_x)_ minima and maxima at each pH value, the midpoint correction provides an unbiased estimate between these limits. Furthermore, experimental pH_mtx v_alues determined from SypHer3 fluorescence ranged from approximately pH 6.4-7.4, over which the correction factors derived from the bound or free occupancy states differed modestly (Fig. 5c). Taken together, uncertainty related to the Ca^2+^ occupancy of R-GECO1 is expected to have a limited impact on the corrected fluorescence signal.

### Statistics and reproducibility

Data are reported as mean ± S.D. unless otherwise stated. A sample size of at least three biological replicates was used for all studies. Statistical analyses were performed in GraphPad Prism 10, and statistical tests used are indicated throughout the text and in figure legends. P values < 0.05 were considered statistically significant.

